# Co-targeting strategy for precise, scarless gene editing with CRISPR/Cas9 and donor ssODNs in *Chlamydomonas*

**DOI:** 10.1101/2021.03.26.437214

**Authors:** Soujanya Akella, Xinrong Ma, Romana Bacova, Zachary P. Harmer, Martina Kolackova, Xiaoxue Wen, David A. Wright, Martin H. Spalding, Donald P. Weeks, Heriberto Cerutti

**Affiliations:** School of Biological Sciences and Center for Plant Science Innovation, University of Nebraska–Lincoln, Lincoln, Nebraska 68588, USA; Department of Chemistry and Biochemistry, Mendel University in Brno, Zemedelska 1, CZ-613 00, Brno, Czech Republic; Department of Genetics, Development and Cell Biology, Iowa State University, Ames, Iowa 50011, USA; Department of Biochemistry, University of Nebraska-Lincoln, Lincoln, Nebraska 68588, USA

**Author notes:** **Corresponding author:** Heriberto Cerutti, School of Biological Sciences and Center for Plant Science Innovation, University of Nebraska-Lincoln, 1901 Vine Street, Lincoln, Nebraska 68588, USA. **Author contributions:** D.A.W., M.H.S., D.P.W. and H.C. conceived and designed the research. S.A., X.M., R.B., Z.P.H., M.K., X.W. and D.A.W. performed the experiments. S.A., X.M., D.A.W., M.H.S., D.P.W. and H.C. analyzed the data. S.A., D.P.W. and H.C. wrote the manuscript. All the authors read and approved the article. These authors contributed equally to this work (Soujanya Akella, Xinrong Ma). State Key Laboratory of Food Nutrition and Safety, College of Biotechnology, Tianjin University of Science & Technology, Tianjin 300457, China (Xinrong Ma). Microbiology and Antimicrobial Resistance Department, Veterinary Research Institute, Hudcova 296/70 Brno, Czech Republic (Romana Bacova). Senior authors (Donald P. Weeks, Heriberto Cerutti).

## Abstract

Programmable site-specific nucleases, such as the CRISPR/Cas9 ribonucleoproteins (RNPs), have allowed creation of valuable knockout mutations and targeted gene modifications in Chlamydomonas. However, in walled strains, present methods for editing genes lacking a selectable phenotype involve co-transfection of RNPs and exogenous double-stranded DNA (dsDNA) encoding a selectable marker gene. Repair of the double-stranded DNA breaks induced by the ribonucleoproteins is usually accompanied by genomic insertion of exogenous dsDNA fragments, hindering the recovery of precise, scarless mutations in target genes of interest. In this study, we tested whether co-targeting two genes by electroporation of pairs of CRISPR/Cas9 RNPs and single-stranded oligodeoxynucleotides (ssODNs) would facilitate the recovery of precise edits in a gene of interest (lacking a selectable phenotype) by selection for precise editing of another gene (creating a selectable marker) - in a process completely lacking exogenous dsDNA. We used *PPX1* (encoding protoporphyrinogen IX oxidase) as the generated selectable marker, conferring resistance to oxyfluorfen, and identified precisely, scarless edited *FTSY* or *WDTC1* genes in ∼1% of the oxyfluorfen resistant colonies. Analysis of the target site sequences in edited mutants suggested that ssODNs were used as templates for DNA synthesis during homology directed repair, a process prone to replicative errors. The Chlamydomonas acetolactate synthase gene could also be efficiently edited to serve as an alternative selectable marker. This transgene-free strategy may allow creation of individual strains containing precise mutations in multiple target genes, to study complex cellular processes, pathways or structures.

**One sentence summary:** Co-targeting two genes by co-electroporation of CRISPR/Cas9 RNPs and ssODN repair templates allows concomitant genome editing to create a selectable marker gene and to introduce precise modifications in another gene of interest.

## Introduction

The green alga *Chlamydomonas reinhardtii* has gained recognition as a model organism for the study of diverse organelles and physiological processes and has also been used in explorative work related to biofuel, nutraceutical, and pharmaceutical recombinant protein production (Rosales-Mendoza et al., 2012; Jinkerson and Jonikas, 2015; Scranton et al., 2015; Salomé and Merchant, 2019). The ability to genetically manipulate the Chlamydomonas genome allows the investigation of gene function as well as the pursuit of innovative biotechnological applications. A variety of methods have been developed to disrupt nuclear genes in this alga, including chemical-, UV light-, gamma irradiation-, and insertional-mutagenesis, the latter involving cell transfection with exogenous DNA that integrates randomly into the genome (Jinkerson and Jonikas, 2015; Picariello et al., 2020). Moreover, a genome-wide mutant library has been recently generated using insertional mutagenesis (Li et al., 2016; 2019). However, all these methods result in random alterations to the genome sequence, and mutations in a desired gene may not be obtained or may not affect gene function.

Targeted strategies for nuclear gene inactivation have also been explored in Chlamydomonas. Post-transcriptional gene silencing by RNA interference (RNAi), involving transgenic strains expressing artificial microRNAs (Molnar et al., 2009; Zhao et al., 2009) or double-stranded RNAs (Rohr et al., 2004; Kim and Cerutti, 2009), seems to have variable knock-down efficiency, depending on the target gene, and may potentially cause unintended off-target effects. Homologous recombination (HR)-mediated gene disruption has also been attempted, but a major drawback is the low frequency of HR between exogenous DNA and a nuclear gene of interest (Sodeinde and Kindle, 1993; Gumpel et al., 1994; Zorin et al., 2009; Sizova et al., 2013; Jinkerson and Jonikas, 2015; Jiang et al., 2017). Promisingly, recent studies have successfully used sequence specific nucleases (SSNs) to achieve targeted gene disruption (Sizova et al., 2013; Jiang et al., 2014; Baek et al., 2016; Shin et al., 2016; Ferenczi et al., 2017; Greiner et al., 2017; Jiang and Weeks, 2017; Shamoto et al., 2018; Guzmán-Zapata et al., 2019; Angstenberger et al., 2020; Cazzaniga et al., 2020; Kang et al., 2020; Kim et al., 2020; Park et al., 2020; Picariello et al., 2020). SSNs cause DNA double-strand breaks (DSBs) at specific sites, which can be repaired by alternative cellular mechanisms resulting in mutations or precise sequence changes. DSB repair by error-prone non-homologous end-joining (NHEJ), including canonical and alternative pathways, may result in insertions/deletions (i.e., indels) and/or missense/nonsense mutations at the target sites (Rodgers and McVey, 2016; Gallagher and Haber, 2017; Capdeville et al., 2020; Gallagher et al., 2020; Scully et al., 2020). Alternatively, homology directed repair (HDR) in the presence of template donor DNA, which can also occur by several pathways, may result in precise sequence changes (Rodgers and McVey, 2016; Gallagher and Haber, 2017; Paix et al., 2017; Capdeville et al., 2020; Gallagher et al., 2020; Scully et al., 2020).

Most current work on nuclear gene targeting in Chlamydomonas has focused on the RNA-programmable site-specific nucleases, such as the clustered regularly interspaced short palindromic repeat (CRISPR)/CRISPR associated protein 9 (Cas9) system from *Streptococcus pyogenes* (Makarova et. al., 2011; Jinek et al., 2012; Jeon et al., 2017; Swarts and Jinek, 2018), which rely on Watson-Crick base-pairing for DNA recognition. Several reports have demonstrated that both CRISPR/Cas9 and CRISPR/Cas12a (formerly Cpf1), belonging to type II-A and V-A of the CRISPR-Cas systems (Makarova et. al., 2011; Swarts and Jinek, 2018), are useful for targeted gene disruption in Chlamydomonas (Jiang et al., 2014; Baek et al., 2016; Shin et al., 2016; Ferenczi et al., 2017; Greiner et al., 2017; Jiang and Weeks, 2017; Shamoto et al., 2018; Guzmán-Zapata et al., 2019; Angstenberger et al., 2020; Cazzaniga et al., 2020; Kang et al., 2020; Kim et al., 2020; Park et al., 2020; Picariello et al., 2020). In this organism, the CRISPR/Cas systems have been implemented as a transgenic method, with components expressed either transiently from plasmids (Jiang et al., 2014; Guzmán-Zapata et al., 2019) or constitutively from genome integrated constructs (Greiner et al., 2017; Jiang and Weeks, 2017; Park et al., 2020), or as a transgene-free approach, by introducing into cells preassembled CRISPR/Cas9 (or CRISPR/Cas12a) ribonucleoproteins (RNPs) by electroporation (Baek et al., 2016; Shin et al., 2016; Ferenczi et al., 2017; Greiner et al., 2017; Shamoto et al., 2018; Angstenberger et al., 2020; Cazzaniga et al., 2020; Kim et al., 2020; Picariello et al., 2020) or by using a cell penetrating peptide (Kang et al., 2020). Published methods using RNA-programmable site-specific nucleases differ in efficiency, applicable genes/strains, and/or experimental details but, in general, targeted disruption of nuclear genes in Chlamydomonas and in the related alga *Volvox carteri* has become a feasible approach (Baek et al., 2016; Shin et al., 2016; Ferenczi et al., 2017; Greiner et al., 2017; Jiang and Weeks, 2017; Shamoto et al., 2018; Guzmán-Zapata et al., 2019; Ortega-Escalante et al., 2019; Angstenberger et al., 2020; Cazzaniga et al., 2020; Kang et al., 2020; Kim et al., 2020; Park et al., 2020; Picariello et al., 2020).

In contrast, precise gene editing (i.e., precise nucleotide changes in a target genomic sequence) triggered by RNA-programmable SSNs remains fairly inefficient or limited to specific strains/experimental approaches (Ferenczi et al., 2017; Greiner et al., 2017; Jiang and Weeks, 2017). As reported for somatic cells of land plants (Capdeville et al., 2020), a major constraint is that, in Chlamydomonas, DSB repair by error-prone NHEJ pathways appears to be much more efficient than homologous recombination in the presence of donor DNA (Sizova et al., 2013; Greiner et al., 2017). As described for some mammalian cell lines (Shy et al., 2016; Mitzelfelt et al., 2017), only a small subset of the population, in asynchronously grown Chlamydomonas, may be capable of HDR, possibly associated with being in a certain phase of the cell cycle (Angstenberger et al., 2020). Additionally, the cell wall appears to pose a significant barrier for the introduction of macromolecules into Chlamydomonas cells (Jeon et al., 2017; this work). Electroporation of CRISPR/Cas RNPs into cell-wall-less mutant strains or after removal of the cell wall by autolysin treatment often resulted in gene editing (disruption) frequencies of a few percentage (relative to the number of electroporated cells) (Baek et al., 2016; Ferenczi et al., 2017; Shamoto et al., 2018; Angstenberger et al., 2020; Picariello et al., 2020). However, using similar methods with walled strains resulted in gene editing (disruption) frequencies several orders of magnitude lower (Shin et al., 2016; Greiner et al., 2017; Kang et al., 2020; this work). In the absence of a selectable phenotype for a gene of interest, a common approach has been to electroporate walled strains with CRISPR/Cas RNPs and DNA coding for a selectable marker (e.g., an antibiotic resistance gene), sometimes with the addition of template DNA to elicit HDR. Chlamydomonas colonies surviving on the selective agent, due to nuclear integration and expression of the selectable transgene, were then screened for editing of a desired endogenous gene in labor-intensive efforts (Shin et al., 2016; Greiner et al., 2017; this work). Unfortunately, in a high percentage of cases, the target gene contained insertions of the exogenous marker DNA or other DNA fragments, preventing the recovery of precisely edited genes for study (Greiner et al., 2017; this work).

To overcome the outlined problems, we explored whether selection for precise editing of an endogenous gene in Chlamydomonas, which would allow exclusive selection of cells taking up the editing components and capable of carrying out HDR, may facilitate the recovery of precise, scarless edits in another gene of interest, when both genes were simultaneously targeted by co-electroporation of CRISPR/Cas9 RNPs and template donor DNAs. We used single-stranded oligodeoxynucleotides (ssODNs) as donor DNA because prior work demonstrated their usefulness as templates in the repair of CRISPR/Cas-induced DSBs in Chlamydomonas (Ferenczi et al., 2017; Greiner et al., 2017; Jiang and Weeks, 2017) and land plants (Shan et al., 2013; Svitashev et al., 2015; Sauer et al., 2016; Yi and Goshima, 2020).

The enzyme protoporphyrinogen oxidase (Protox), encoded by the *PPX1* gene in Chlamydomonas, oxidizes protoporphyrinogen IX to protoporphyrin IX in the biosynthetic pathway of heme and chlorophyll (Duke et al., 1991; Randolph-Anderson et al., 1998). Inhibition of this enzyme causes accumulation of protoporphyrinogen IX, which is non-enzymatically oxidized to protoporphyrin IX and eventually leads to membrane peroxidation and cell lethality in the light (Duke et al., 1991; Ha et al., 2004). A single base pair mutation (G->A, causing a valine-389 to methionine substitution) within the protein coding sequence results in the Protox enzyme being resistant to inhibition by porphyric herbicides such as oxyfluorfen (Randolph-Anderson et al., 1998; Brueggeman et al., 2014). Electroporation of Chlamydomonas cells with a CRISPR/Cas9 (*PPX1*) RNP and a ssODN donor (designed to introduce the G->A mutation in *PPX1* during HDR) led to the isolation of colonies precisely edited in *PPX1* by selection on oxyfluorfen containing medium. Interestingly, co-targeting *PPX1* (as the selectable marker) and *FTSY* (encoding a signal recognition particle receptor protein which is required for the integration of light-harvesting complex proteins into thylakoid membranes) or *WDTC1* (encoding a conserved protein with antiadipogenic functions in several eukaryotes) allowed the recovery of precisely edited mutants in the latter genes. Optimizing this approach of simultaneously editing a selectable marker and any gene of interest may prove to be a viable strategy for introducing precise sequence changes in the genome of walled Chlamydomonas and, potentially, of other microalgae.

## Results

### Targeted disruption of the *FTSY* gene

The *FTSY* gene encodes a component of the chloroplast signal recognition particle-dependent pathway, which is required for insertion of light-harvesting chlorophyll a/b-binding proteins into the thylakoid membranes (Aldridge et al., 2009; Kirst et al., 2012). In Chlamydomonas, null *FTSY* mutants showed diminished ability to assemble some proteins of the light harvesting complex II, resulting in lower chlorophyll content than in the wild type (Kirst et al., 2012; Baek et al., 2016; Kim et al., 2020). Hence, mutant colonies displayed a pale-green phenotype. Because of this easily identifiable phenotype, we initially attempted to edit exon 4 of the *FTSY* gene (Fig. 1A), using a CRISPR/Cas9 protocol dependent on square-wave electroporation as described by Greiner et al. (2017).

**Figure 1.**
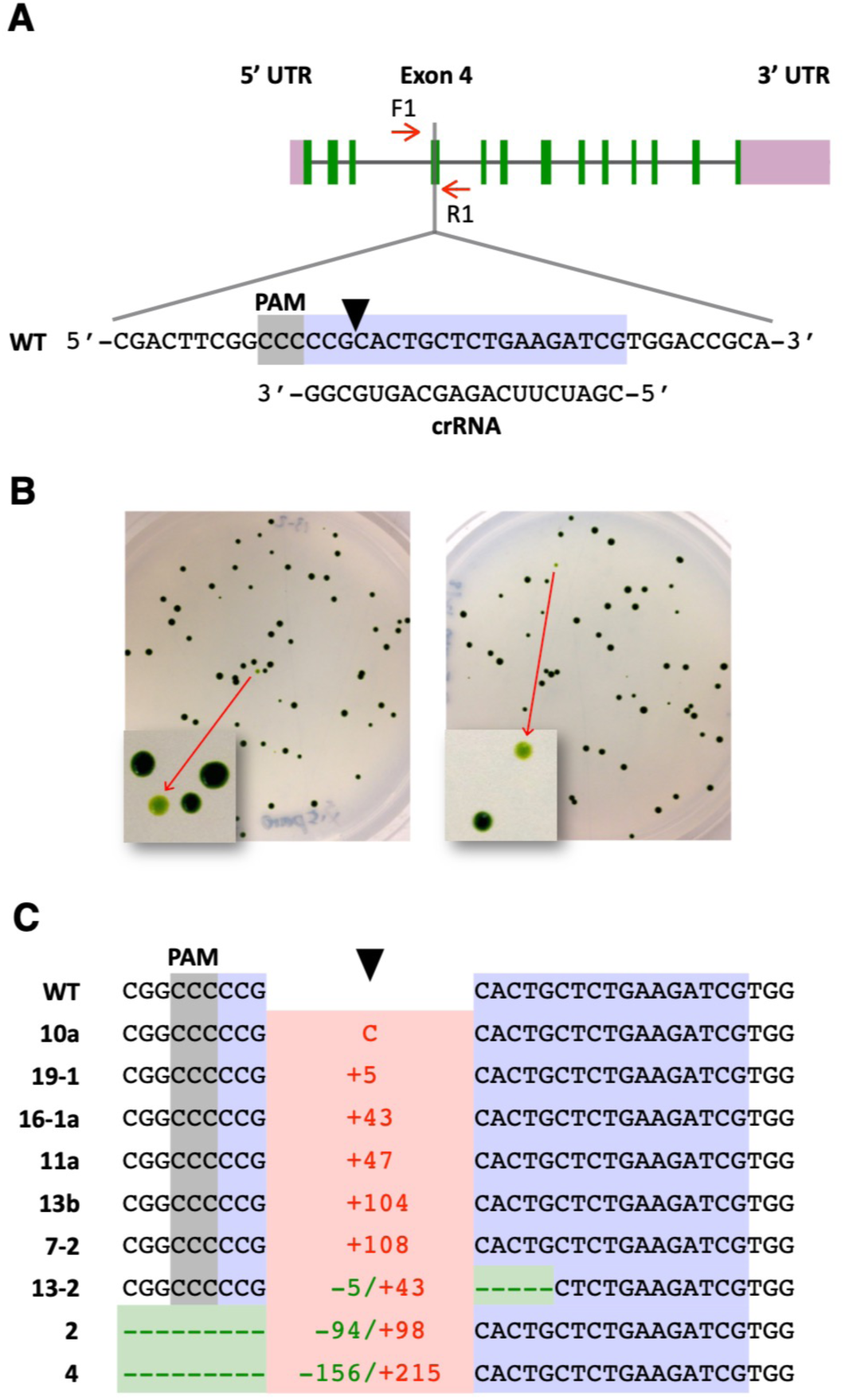
Targeted disruption of the *FTSY* gene. A, Schematic of the *FTSY* gene showing the target (highlighted in bluish purple) and PAM (highlighted in gray) sequences. A black arrowhead indicates the Cas9 cleavage site. Short red arrows indicate the primers used for PCR analyses. B, Cells co-transfected with the CRISPR/Cas9 (*FTSY*) RNP and dsDNA encoding the *aphVIII* transgene were spread on TAP agar plates containing paromomycin. Representative plates show some pale green colonies as a consequence of *FTSY* gene disruption. C, DNA sequences of pale green colonies, indicating alterations at the *FTSY* target site relative to the wild type (WT). Insertions, indicating type (C, cytosine) or number of base pairs, are depicted in red. Deletions, indicating type or number of base pairs, are depicted in green. Complete sequences are shown in Supplemental Fig. 2.

Cells of the walled CC-5415 (g1) strain were electroporated with a CRISPR/Cas9 (*FTSY*) RNP and a dsDNA PCR fragment containing a transgene expressing the *Streptomyces rimosus aphVIII* gene, which confers resistance to the antibiotic paromomycin (Sizova et al., 2001). After electroporation, cells were selected on agar plates containing paromomycin. Antibiotic resistant colonies displaying a pale green phenotype (Fig. 1B) were assumed to be *FTSY* null mutants for calculation of gene disruption frequencies (Supplemental Table 1). However, we note that the reported values are an approximation to the actual frequency of targeted gene alterations since some CRISPR/Cas9-induced mutations in *FTSY* may not affect protein function and the corresponding colonies would not be detectable as pale green. Conversely, some pale green colonies may be caused by random insertion of the selectable transgene into genes, other than *FTSY*, associated with chlorophyll synthesis or photosynthetic complex assembly. Yet, all pale green colonies examined by PCR amplification of the target site (Supplemental Table 1) either showed indels at the Cas9 cleavage site (Fig. 1C), consistent with disruption of *FTSY* gene function, or lacked a detectable PCR product, likely due to large DNA insertions and/or rearrangements at the target site, a previously reported common occurrence associated with CRISPR/Cas9 gene editing in Chlamydomonas (Shin et al., 2016; Greiner et al., 2017; Kang et al., 2020; Picariello et al., 2020).

In agreement with the results of Greiner et al. (2017), heat shocking the cells prior to electroporation increased >5-fold the number of recovered pale green colonies (Supplemental Table 1, Experiments 2 and 3). To achieve precise *FTSY* gene editing, in several experiments, we also added to the electroporation mix a ssODN (Supplemental Table 2), overlapping the Cas9 cleavage site and designed to serve as template for HDR. Precise repair by homology directed mechanisms of the DSB caused by CRISPR/Cas9 (*FTSY*) RNP would introduce stop codons within the *FTSY* coding sequence, create a new restriction enzyme site (*Nhe*I) for genotypic analyses, and destroy the Cas9 protospacer adjacent motif (PAM) (Supplemental Fig. 1). However, out of 21 pale green colonies examined by PCR amplification of the targeted *FTSY* exon, 12 lacked a detectable PCR product and 9 showed indels at the target site (Fig. 1C and Supplemental Fig. 2). We did not recover any colony precisely edited by HDR in any of these experiments, underlining the difficulty in obtaining precise gene editing in walled Chlamydomonas strains with current methodological approaches. Although, one colony exhibited apparent integration of donor DNA by homologous recombination on one side of the DSB whereas the repair was carried out by NHEJ on the other side, resulting in the insertion of a 104-bp DNA fragment (Supplemental Fig. 2, colony 13b) (see Discussion).

**Figure 2.**
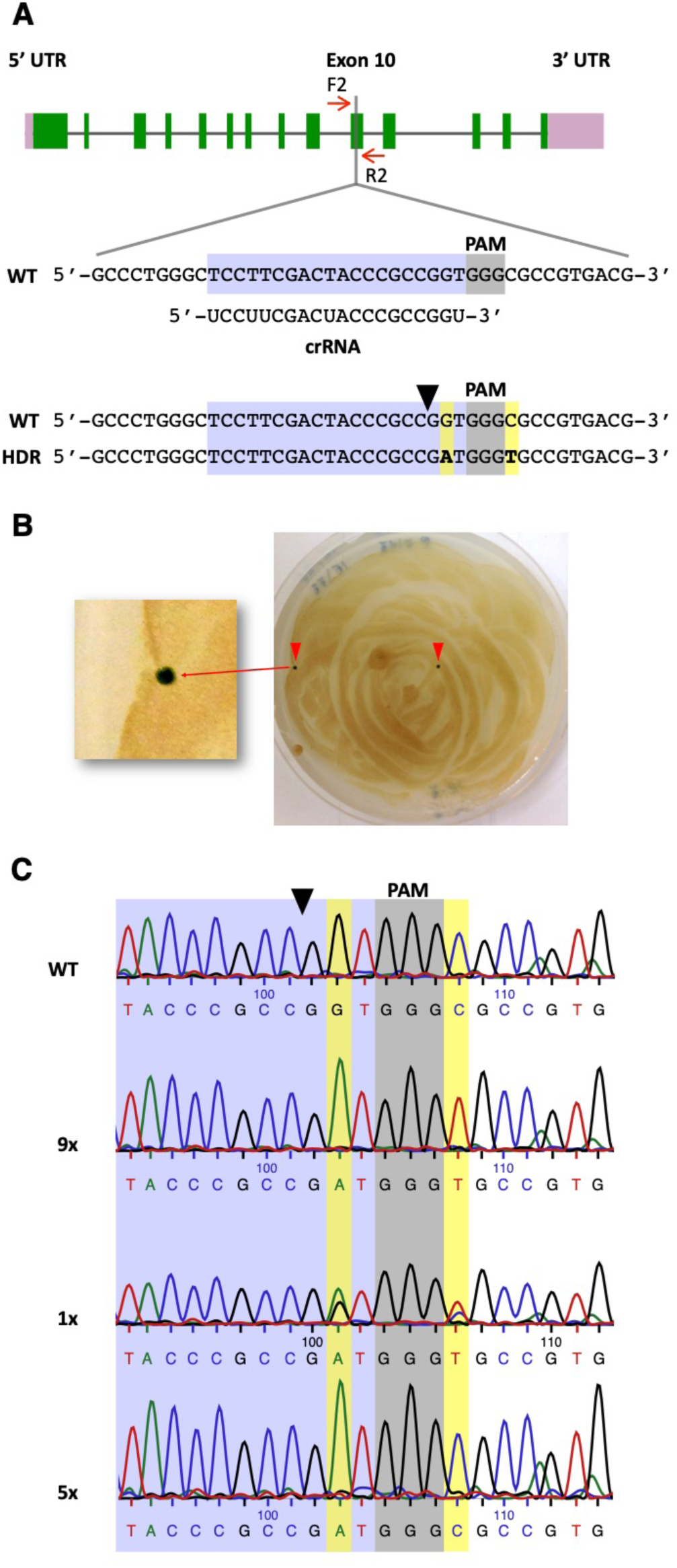
Precise editing of the *PPX1* gene. A, Schematic of the *PPX1* target region in exon 10. Color schemes and symbols are as described under Fig. 1A. Homology directed repair (HDR) of the DSB, using as template the transfected *PPX1* ssODN, is expected to introduce two base pair changes (highlighted in yellow) into the genome (bottom sequence). B, Cells co-transfected with the CRISPR/Cas9 (*PPX1*) RNP and ssODN donor DNA were spread on TAP agar plates containing oxyfluorfen. A representative plate shows two green oxyfluorfen resistant colonies as a consequence of *PPX1* gene editing. C, Representative sequencing chromatograms of the wild type (WT), fully edited (9x, containing both G->A and C->T changes) and, likely, partly edited (5x, containing only the G->A change) colonies. The chromatogram of an apparently mixed colony (1x), displaying a heterozygous (i.e., wild-type and fully edited) sequence, is also shown. Figures followed by an X indicate the number of colonies of each DNA type examined by sequencing.

We also examined uptake of the CRISPR/Cas9 (*FTSY*) RNP after electroporation of the walled g1 strain of Chlamydomonas. For these experiments, we assembled the RNP with trans-activating CRISPR RNA (tracrRNA) conjugated to the ATTO 550 fluorophore (Banas et al., 2020). Four hours after electroporation, cells were examined by fluorescence microscopy to visualize uptake and intracellular localization of the labeled RNP. The tracrRNA-ATTO 550 electroporated alone localized predominantly in the nucleus (Supplemental Fig. 3A), as previously reported for single-stranded DNA oligonucleotides (Jiang et al., 2017). In contrast, the assembled tracrRNA-ATTO 550-Cas9 RNP was distributed in the cytosol and the nucleus with some preferential perinuclear accumulation (Supplemental Fig. 3A). However, examining over a thousand individual cell images revealed that less than 1% of the cells had taken up enough RNP to be detectable by fluorescence microscopy (Supplemental Fig. 3B), suggesting that electroporation of CRISPR/Cas9 RNPs into walled Chlamydomonas strains is quite inefficient (at least under our experimental conditions).

### Precise editing of the *PPX1* gene

We next attempted to precisely modify an endogenous gene amenable to selection, with the goal of developing a co-editing approach in Chlamydomonas. As already mentioned, a single point mutation (G->A) in *PPX1* exon 10 (Fig. 2A), causing a single amino acid substitution (Val389Met) in the Protox enzyme, confers resistance to porphyric herbicides in Chlamydomonas (Randolph-Anderson et al., 1998; Brueggeman et al., 2014). Cells of the g1 strain were electroporated with a CRISPR/Cas9 (*PPX1*) RNP and a ssODN (Supplemental Table 2), overlapping the Cas9 cleavage site and designed to serve as template for HDR. Precise DSB repair by homology directed mechanisms would introduce the G->A mutation within the *PPX1* coding sequence as well as a nearby, functionally silent, C->T mutation (Fig. 2A and Supplemental Fig. 4). Electroporated cells were selected on agar plates containing oxyfluorfen. By using the protocol developed by Greiner et al. (2017), we recovered several oxyfluorfen resistant colonies (Fig. 2B). Moreover, the frequency of CRISPR/Cas9 (*PPX1*) RNP induced mutants, particularly when including a heat shock treatment (Supplemental Table 3, Experiments 1 and 2), was well above the frequency of spontaneous *PPX1* mutation to herbicide resistance, which has been reported to be <1×10^−8^ (Randolph-Anderson et al., 1998). However, the overall number of oxyfluorfen resistant colonies was quite low.

To enhance the recovery of *PPX1* edited colonies we introduced several modifications to the Greiner et al. (2017) method, including changes to some electroporation parameters, the amounts of Cas9 nuclease and ssODN donor used in the electroporation mix, as well as the timing of the heat shock treatment, which was performed after (rather than before) cell electroporation. All these modifications have been incorporated into our optimized protocol, described under Materials and Methods. The modified protocol increased ∼5-fold the number of recovered oxyfluorfen resistant colonies (Supplemental Table 3, Experiments 3 and 4). Fifteen of these colonies were examined by PCR amplification of the target site and sequencing of the PCR products. Nine colonies displayed the expected (i.e., G->A and C->T) sequence changes, one colony appeared to be a mixture between wild type and a properly edited clone, and five colonies showed only the G->A change, which is solely necessary to confer herbicide resistance (Fig. 2C). Thus, about two thirds of the examined colonies were fully consistent with editing by HDR, using as template the electroporated ssODN donor. However, we surmise that the remaining examined colonies were also edited by homology directed mechanisms, since we have not recovered any spontaneous herbicide resistant mutant in any of our negative control experiments (Supplemental Table 3). As described in other eukaryotes (Gallagher and Haber, 2017; Paix et al., 2017; Boel et al., 2018; Gallagher et al., 2020), we expected that Cas9-induced DSBs in Chlamydomonas would be repaired by the single-strand template repair (SSTR) mechanism (see Discussion). Following this model (Supplemental Fig. 4), the *PPX1* ssODN would be used as a homologous template for DNA synthesis primed by the 3’ ending strand on the left side of the cleavage site. If a very short stretch of DNA (at most 6 nucleotides), next to the Cas9 cleavage site, is copied by DNA synthesis, only the G->A change would be introduced into the genome of some cells. Alternatively, a longer stretch of DNA may be copied (including both G->A and C->T modifications) but after annealing of the newly synthesized strand with the complementary wild-type strand, heteroduplex DNA may be formed and mismatch repair mechanisms may correct the C->T change back to the original sequence (Harmsen et al., 2018; Gallagher et al., 2020). Either of these alternative modes of DNA repair would explain the recovery of colonies showing only the G->A edit.

### Precise co-editing of the *PPX1* and *FTSY* genes

If in asynchronously grown Chlamydomonas only a small subpopulation of cells is receptive to DSB repair by homology directed mechanisms, selection for an HDR-based editing event (such as *PPX1* editing) may facilitate the recovery of simultaneous HDR-mediated edits at other loci. To test this hypothesis, we attempted co-editing of *PPX1* and *FTSY* by co-electroporation of the respective CRISPR/Cas9 RNPs and template ssODN donors. Electroporated cells were selected on agar plates containing oxyfluorfen. Out of three independent experiments, we isolated 9 (4.0%) pale green colonies, with alterations in the *FTSY* gene, among 223 oxyfluorfen resistant colonies (Supplemental Table 4). However, only 3 (1.3%) of these colonies displayed changes in the *FTSY* sequence (Fig. 3A) consistent with precise, scarless editing by HDR. In these cases, a new *Nhe*I restriction enzyme site, as shown for a subset of these colonies (Fig. 3B), and the designed stop codons within the *FTSY* coding sequence were incorporated at the intended site (Fig. 3C, colonies 10-1b, 8-1, and 6-1). The other 6 colonies showed indels at or near the expected Cas9 cleavage site (Fig. 3C and Supplemental Fig. 5), although *FTSY* in one of these colonies did appear to be repaired by homology directed mechanisms but including an unexpected single base pair deletion (Fig. 3C, colony 3-2), possibly resulting from replicative errors (see Discussion).

**Figure 3.**
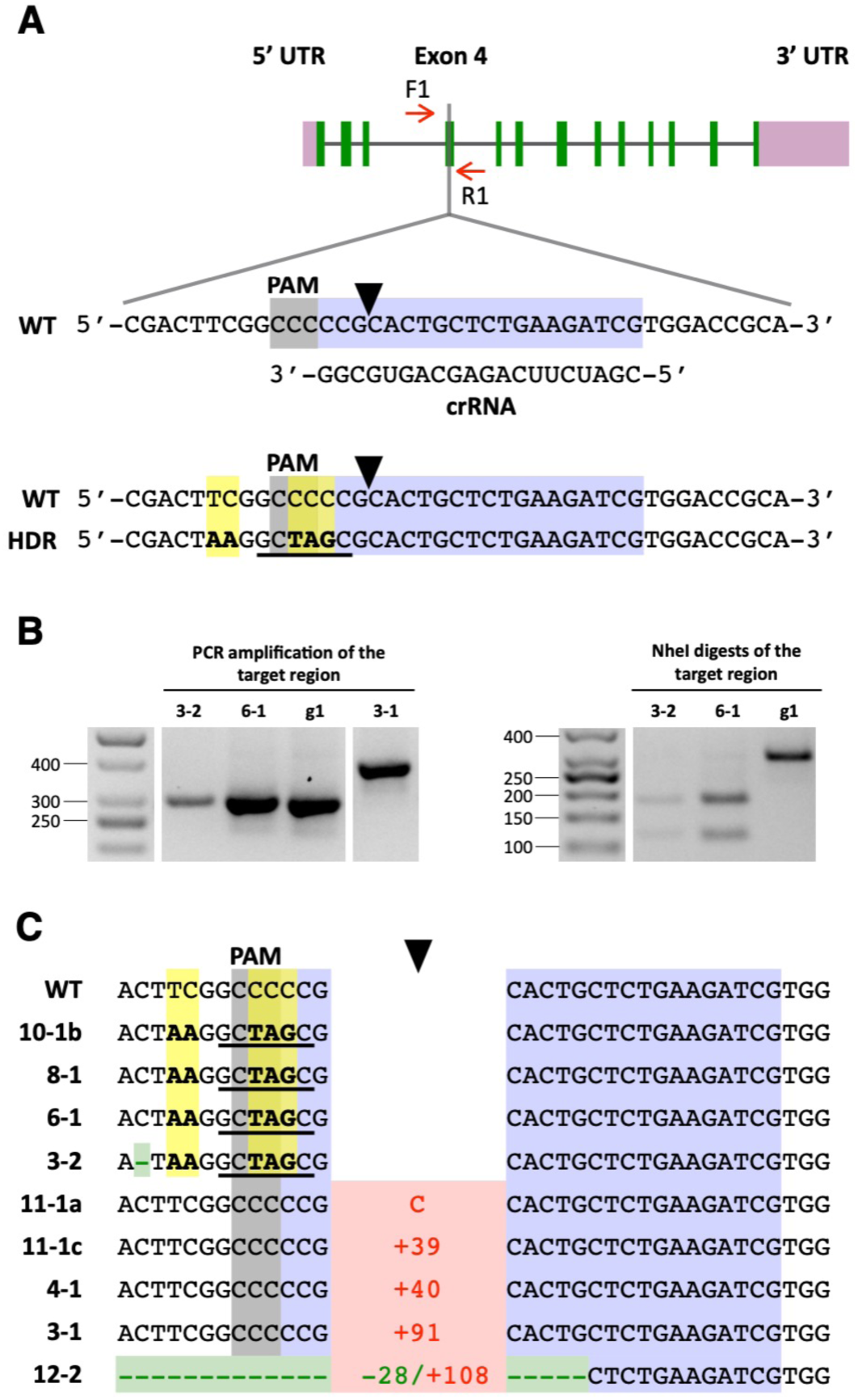
Co-editing of the *FTSY* gene in colonies selected for precise *PPX1* editing. A, Schematic of the *FTSY* target region in exon 4. Color schemes and symbols are as described under Fig. 1A. Homology directed repair (HDR) of the DSB, using as template the transfected *FTSY* ssODN, is expected to introduce five base pair changes (highlighted in yellow) into the genome (bottom sequence). These sequence changes insert two stop codons (i.e., TAA and TAG) in the coding frame, destroy the PAM site, and create a new *Nhe*I restriction enzyme site (underlined in black). B, The *FTSY* target region of selected oxyfluorfen resistant, pale-green colonies was amplified by PCR (with primers F1 and R1) and the PCR products digested with *Nhe*I. The panels show representative reverse images of agarose resolved PCR products stained with ethidium bromide. The sizes of molecular weight markers are indicated in base pairs. g1, wild-type strain. C, DNA sequences of oxyfluorfen resistant, pale green colonies, indicating alterations at the *FTSY* target site relative to the wild type (WT). Insertions, indicating type (C, cytosine) or number of base pairs, are depicted in red. Deletions, indicating type or number of base pairs, are depicted in green. Base substitutions are highlighted in yellow. Complete sequences for colonies exhibiting *FTSY* indels are shown in Supplemental Fig. 5.

As discussed above, in our initial attempts at precise *FTSY* editing, by supplying a ssODN as template for HDR together with dsDNA encoding a selectable marker, out of 21 examined pale green colonies, none showed precise HDR. In contrast, when selecting first for *PPX1* edited colonies on medium containing oxyfluorfen, out of 9 examined pale green colonies, 3 showed precise *FTSY* HDR and an additional one showed *FTSY* sequence changes consistent with repair by homology directed mechanisms, although including an unintended single base pair deletion. Thus, in comparison to current precise gene editing methodology for walled Chlamydomonas strains, our observations suggest that a CRISPR/Cas9 RNP co-targeting strategy does facilitate the isolation of colonies precisely edited in a gene of interest lacking a selectable phenotype.

### Precise co-editing of the *PPX1* and *WDTC1* genes

WD and Tetratricopeptide Repeats 1 (WDTC1) is a conserved eukaryotic protein, containing WD40 and tetratricopeptide repeat domains. In flies and mammals, loss of function of *WDTC1* results in an increase in adipocytes, fat accumulation, and obesity (Häder et al., 2003; Groh et al., 2016). The *Arabidopsis thaliana* homolog, ASG2, also appears to have antiadipogenic functions, since *asg2* knockout mutants produce seeds that have greater weight and increased fatty acid content than the wild type (Ducos et al., 2017). The Chlamydomonas *WDTC1* gene has not been characterized in any detail and the phenotype(s), if any, of a null mutant is unknown. Therefore, it provided a good system for testing the practicality of our co-targeting approach, because edited colonies in the gene of interest needed identification by bulk PCR analyses.

Cells of the g1 strain were co-electroporated with CRISPR/Cas9 RNPs and template ssODN donors targeting *PPX1* and *WDTC1*. Electroporated cells were then selected on agar plates containing oxyfluorfen. Precise repair of the DSB caused by CRISPR/Cas9 (*WDTC1*) RNP, using a ssODN (Supplemental Table 2) as a homologous template, would eliminate the *WDTC1* start codon (precluding synthesis of a full length protein) and create a new *Bss*HII restriction enzyme site for genotypic analyses (Fig. 4A and Supplemental Fig. 6A). Out of two independent experiments, we isolated 9 (2.3%) colonies with alterations in the *WDTC1* gene, among 395 oxyfluorfen resistant colonies examined by PCR (Supplemental Table 5). However, only 3 (0.8%) of these colonies displayed changes in the *WDTC1* sequence (Fig. 4A) consistent with precise, scarless editing by HDR. In these cases, the new *Bss*HII restriction enzyme site was incorporated into the genome, as shown for a subset of these colonies (Fig. 4B), and the *WDTC1* sequence around the start codon (i.e., AATG) was replaced with the intended sequence (i.e., GCGC) (Fig. 4C, colonies 3-15b, 1A-37, and 3B-48). In one colony, all changes expected for HDR did occur but we also observed unintended substitution of two base pairs (Fig. 4C, colony 2B-19), possibly the result of errors during DNA synthesis (see Discussion). The remaining 5 colonies showed indels at the expected Cas9 cleavage site (Fig. 4C and Supplemental Fig. 7).

**Figure 4.**
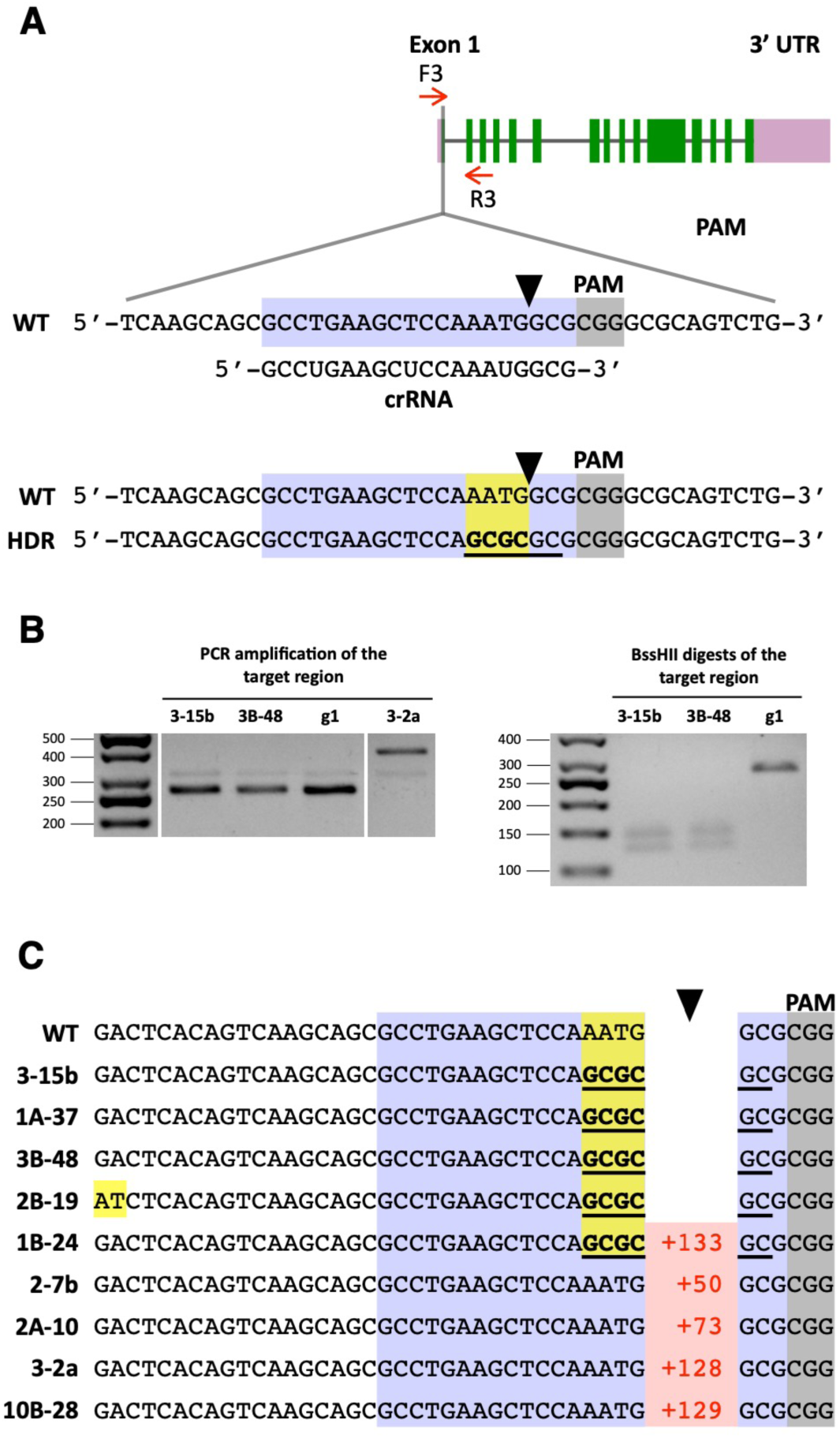
Co-editing of the *WDTC1* gene in colonies selected for precise *PPX1* editing. A, Schematic of the *WDTC1* target region in exon 1. Color schemes and symbols are as described under Fig. 1A. Homology directed repair (HDR) of the DSB, using as template the transfected *WDTC1* ssODN, is expected to introduce four base pair changes (highlighted in yellow) into the genome (bottom sequence). These sequence changes destroy the *WDTC1* start codon and create a new *Bss*HII restriction enzyme site (underlined in black). B, The *WDTC1* target region of selected oxyfluorfen resistant colonies was amplified by PCR (with primers F3 and R3) and the PCR products digested with *BssH*II. The panels show representative reverse images of agarose resolved PCR products stained with ethidium bromide. The sizes of molecular weight markers are indicated in base pairs. g1, wild-type strain. C, DNA sequences of oxyfluorfen resistant colonies showing alterations at the *WDTC1* target site relative to the wild type (WT). Insertions, indicating number of base pairs, are depicted in red. Base substitutions are highlighted in yellow. Complete sequences for colonies exhibiting *WDTC1* indels are shown in Supplemental Fig. 7.

For comparison purposes, we also attempted precise *WDTC1* editing by electroporating g1 cells with CRISPR/Cas9 (*WDTC1*) RNP, a dsDNA PCR fragment containing the *aphVIII* transgene and the *WDTC1* ssODN. Cells were selected on paromomycin containing medium. Out of 96 paromomycin resistant colonies examined by PCR, two (2.1%) showed alterations in the *WDTC1* gene. However, both colonies had insertions of the *aphVIII* transgene at the Cas9 cleavage site, consistent with DSB repair by NHEJ mechanisms (Supplemental Fig. 8, colonies 20 and 63). Although, in one case SSTR appears to have started correctly, using the *WDTC1* ssODN as a homologous repair template (see Discussion), but the event was resolved by NHEJ on the left side of the cleavage site (Supplemental Fig. 8, colony 63). Thus, as for the *FTSY* gene, we did not recover any colony showing precise HDR-mediated editing of *WDTC1* by using established gene editing methodology for walled Chlamydomonas strains.

### Phenotypic characterization of *WDTC1* edited mutants

To gain initial insights on WDTC1 function in Chlamydomonas, we examined growth and the accumulation of nonpolar lipids in a subset of edited mutants. We chose to analyze two precise HDR edited mutants (i.e., 3-15b and 3B-48) and three insertional mutants (i.e., 3-2a, 20 and 63). Semi-quantitative RT-PCR analyses of exon-exon junctions close to the beginning (i.e., exon2/exon3) or to the end (i.e., exon 13/exon14) of the *WDTC1* coding sequence (Fig. 5A) revealed that the HDR edited mutants had similar or slightly higher *WDTC1* transcript abundance than the wild type, when cells were grown under either mixotrophic or phototrophic conditions (Figs. 5B and 5C). In contrast, the insertional mutants displayed reduced *WDTC1* transcript abundance, albeit to different and somewhat variable degrees, relative to the wild type (Figs. 5B and 5C).

**Figure 5.**
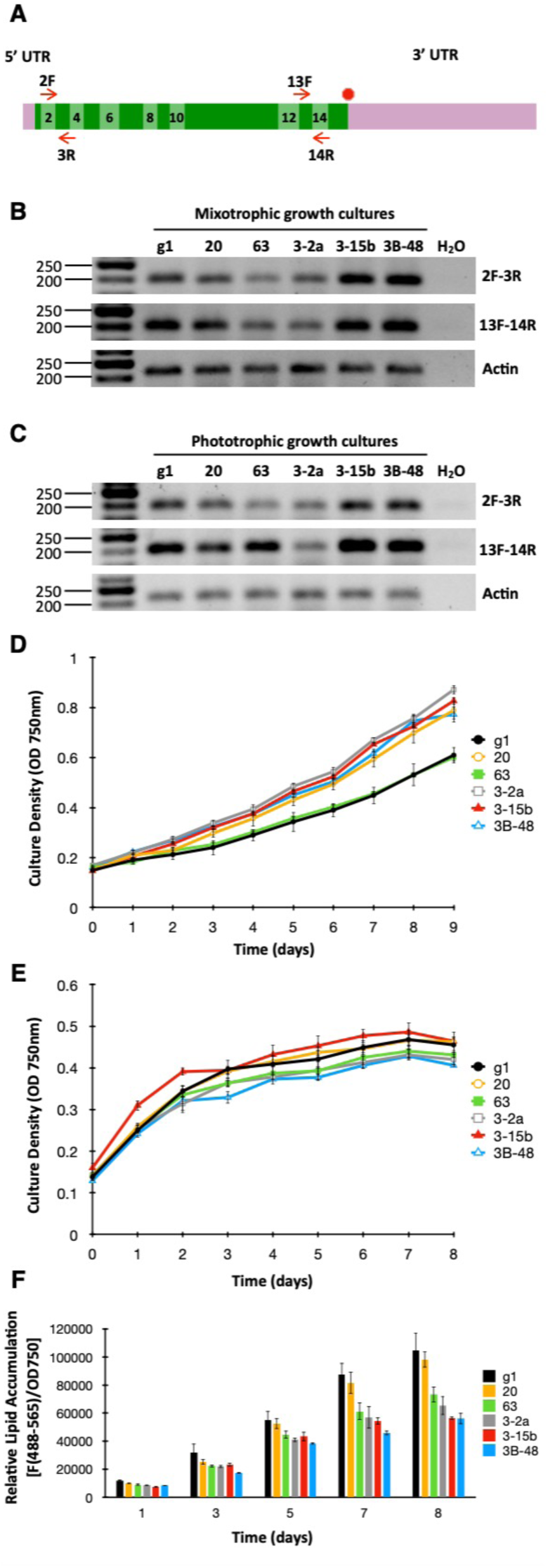
Phenotypic characterization of *WDTC1* mutants generated by CRISPR/Cas9 editing. A, Schematic of the *WDTC1* transcript. Exons are depicted as boxes in two alternating shades of green and even exons are numbered. Short red arrows indicate the primers used for semi-quantitative RT-PCR analyses. B, *WDTC1* transcript abundance examined by semiquantitative RT-PCR in cells grown under mixotrophic conditions. The panels show representative reverse images of agarose resolved RT-PCR products stained with ethidium bromide. Amplification of the Actin transcript was used for normalization purposes. The sizes of molecular weight markers are indicated in base pairs. g1, wild-type strain. C, *WDTC1* transcript abundance examined by semiquantitative RT-PCR in cells grown under phototrophic conditions. D, Growth of the indicated strains under phototrophic conditions in nutrient replete minimal medium. Values shown are the average of three biological replicates ± SD. E, Growth of the indicated strains under mixotrophic conditions in nitrogen deprived medium (TAP-N). Values shown are the average of three biological replicates ± SD. F, Nonpolar lipid accumulation in the indicated strains cultured under mixotrophic conditions in TAP medium lacking nitrogen. Nonpolar lipid content was estimated by staining with the lipophilic fluorophore Nile Red and measuring fluorescence (excitation at 488 nm; emission at 565 nm) in a multiwell plate reader. Nile Red fluorescence was normalized to cell density (determined as absorbance at 750 nm) and expressed in arbitrary units. Values shown are the average of three biological replicates ± SD.

Interestingly, under phototrophic conditions, four of the edited mutants appeared to grow at a somewhat faster rate than the wild type (Fig. 5D), whereas the growth of insertional mutant 63 was nearly identical to that of the wild type. Since WDTC1 homologs have been implicated in antiadipogenic functions in several eukaryotes (Häder et al., 2003; Groh et al., 2016; Ducos et al., 2017), we also examined the performance of the Chlamydomonas mutants with respect to triacylglycerol biosynthesis. This alga has been shown to accumulate significant amounts of triacylglycerol when subject to nitrogen deprivation (Siaut et al., 2011; Msanne et al., 2012; Kim et al., 2018). Thus, we first analyzed the growth/survival of the strains in the absence of nitrogen (in medium supplemented with acetate as a carbon source). Under these conditions, none of the edited mutants differed substantially in growth from the wild type (Fig. 5E). To evaluate neutral lipid accumulation, Chlamydomonas cells were examined by fluorometry after staining with the nonpolar lipid fluorophore Nile Red (Msanne et al., 2012). As expected, the wild-type strain showed substantial accumulation of nonpolar lipids, over time, under nitrogen starvation in the presence of acetate (Fig. 5F). However, four of the Chlamydomonas *WDTC1* edited mutants showed reduced nonpolar lipid accumulation relative to the wild type (Fig. 5F); only insertional mutant 20 behaved similarly to the wild-type strain.

Given that an antiadipogenic role of WDTC1 was not supported by our experiments, defining the function(s) of WDTC1 in Chlamydomonas will require further work. However, in the context of CRISPR/Cas9-generated mutants, we observed that the phenotypes of the two HDR edited mutants (i.e., 3-15b and 3B-48) and of the insertional mutant isolated during co-editing with *PPX1* (i.e., 3-2a) were consistently similar in all the analyses that we performed. In contrast, the *aphVIII* insertional mutants (i.e., 20 and 63) showed divergent behavior under certain conditions. For instance, in comparison to the other edited mutants, mutant 63 differed in growth under phototrophic conditions whereas mutant 20 differed in nonpolar lipid accumulation under nitrogen deprivation. Since the *WDTC1* start codon or the proper translation frame were disrupted in all five examined mutants, these idiosyncratic phenotypic variations cannot be explained based exclusively on loss-of-function of the *WDTC1* gene. The actual reason(s) for the discrepancies is currently unknown. However, these observations raise a cautionary note on the need to carefully examine CRISPR/Cas9-generated insertional mutants since it is conceivable that the unintended integration of exogenous DNA into the Chlamydomonas genome (for instance at off-target sites) may cause phenotypic alterations.

### Precise editing of the *ALS1* gene, an alternative selectable marker

For studies of groups of genes involved in specific enzymatic pathways or cellular functions, the ability to sequentially produce in the same strain (i.e., in the same genetic background) precise mutations in multiple genes of interest can be a powerful tool. However, with the strategy we put forward in the present communication, each new round of mutagenesis with a CRISPR/Cas9 RNP and a ssODN donor targeting a specific gene would require the simultaneous targeting of a different gene that is capable of being converted into a selectable marker. We have already shown that the acetolactate synthase (*ALS1*) gene is an attractive target for CRISPR/Cas9 RNP-directed mutagenesis since a single amino acid change from lysine 257 to threonine (caused by a single base pair substitution, changing codon A**A**G to A**C**G) results in strong resistance to the herbicide sulfometuron methyl (SMM) (Kovar et al., 2002; Jiang and Weeks, 2017). To test the efficiency of *ALS1* as a selectable marker gene as well as an alternative transfection protocol relying on exponential-wave electroporation, cells of the walled CC-124 strain were electroporated with a CRISPR/Cas9 (*ALS1*) RNP and a ssODN (Supplemental Table 2), overlapping the Cas9 cleavage site and designed to serve as template for HDR. Precise DSB repair by homology directed mechanisms would introduce the A->C mutation within the *ALS1* coding sequence as well as a nearby, functionally silent, C->T mutation that creates a new *Eco*RV restriction enzyme site for genotypic analyses (Fig. 6A and Supplemental Fig. 9). Electroporated cells were selected on agar plates containing SMM.

**Figure 6.**
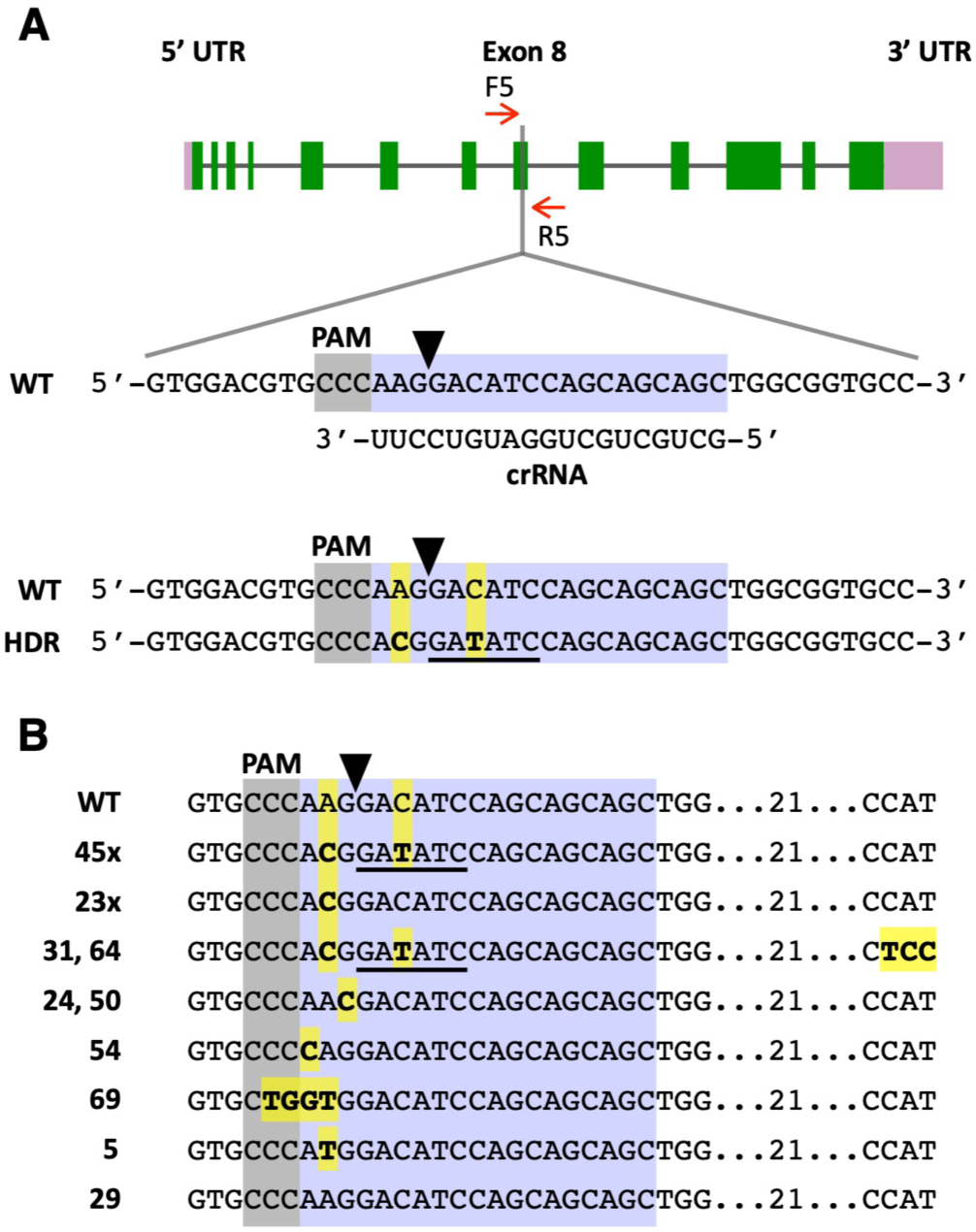
Precise editing of the *ALS1* gene. A, Schematic of the *ALS1* target region in exon 8. Color schemes and symbols are as described under Fig. 1A. Homology directed repair (HDR) of the DSB, using as template the transfected *ALS1* ssODN, is expected to introduce two base pair changes (highlighted in yellow) into the genome (bottom sequence). These sequence changes cause an amino acid substitution (K257T) in the encoded acetolactate synthase enzyme and create a new *Eco*RV restriction enzyme site (underlined in black). B, DNA sequences of sulfometuron methyl resistant colonies showing alterations at the *ALS1* target site relative to the wild type (WT). Base substitutions are highlighted in yellow. Figures followed by an X (i.e., 45x and 23x) indicate the number of colonies of each DNA type analyzed by sequencing (complete sequences for each colony are shown in Supplemental Fig. 10). Other numbers indicate the colony names.

Out of 483 SMM resistant colonies obtained in the treatment with CRISPR/Cas9 (*ALS1*) RNP and ssODN donor, 76 were examined by PCR amplification of the target site and sequencing of the PCR products (Supplemental Table 6). Forty-seven (61.8%) colonies displayed the expected (i.e., A->C and C->T) sequence changes, although in two cases (i.e., colonies 31 and 64) with additional sequence alterations (Fig. 6B and Supplemental Fig. 10A; Supplemental Table 6). Twenty-three (30.2%) colonies only showed the A->C change, which is necessary to confer herbicide resistance (Fig. 6B and Supplemental Fig. 10B; Supplemental Table 6). These results are similar to those obtained in experiments with the *PPX1* gene (see above) and suggest that ∼92% of the SMM resistant colonies are consistent with editing by HDR (Supplemental Table 6), using as homologous template the electroporated ssODN donor. Five additional colonies showed mutations causing substitution of lysine 257 for alternative amino acids (other than threonine) (Fig. 6B, colonies 24, 50, 54, 69 and 5; Supplemental Table 6), which apparently also confer resistance to SMM. In two colonies, a single nucleotide change occurred at the Cas9 cleavage site (Fig. 6B, colonies 24 and 50; Supplemental Table 6), possibly as a result of errors during NHEJ. The other three colonies displayed single or a few nucleotide substitutions close to the cleavage site (Fig. 6B, colonies 54, 69 and 5; Supplemental Table 6) but the mechanism(s) responsible for these changes is not clear. One colony out of the 76 examined had a wild type target sequence (Fig. 6B, colony 29), suggesting a potential spontaneous mutation conferring SMM resistance elsewhere in the *ALS1* gene.

## Discussion

As mentioned earlier, targeted disruption of nuclear genes, mediated by RNA-programmable SSNs, has become a practical method in Chlamydomonas (Baek et al., 2016; Shin et al., 2016; Ferenczi et al., 2017; Greiner et al., 2017; Jiang and Weeks, 2017; Shamoto et al., 2018; Guzmán-Zapata et al., 2019; Angstenberger et al., 2020; Cazzaniga et al., 2020; Kang et al., 2020; Kim et al., 2020; Park et al., 2020; Picariello et al., 2020). In contrast, precise gene editing still occurs at relatively low frequencies or it is limited to specific (cell-wall-less) strains or experimental approaches (Ferenczi et al., 2017; Greiner et al., 2017; Jiang and Weeks, 2017). In Chlamydomonas, as observed in land plants and other eukaryotes (Gallagher and Haber, 2017; Kan et al., 2017; Paix et al., 2017; Boel et al., 2018; Richardson et al., 2018; Sansbury et al., 2019; Capdeville et al., 2020; Gallagher et al., 2020), the repair of DSBs induced by CRISPR/Cas9 RNPs likely occurs by several distinct pathways, partly determined by the repair machinery expressed in each cell, the complexity of the DSB, and the nature of the DNA molecules involved in the repair. As described for some mammalian cell lines (Shy et al., 2016; Mitzelfelt et al., 2017), only a small subset of the population, in asynchronously grown Chlamydomonas, may be capable of HDR, possibly associated with being in a certain phase of the cell cycle (Angstenberger et al., 2020). In addition, the cell wall appears to pose a significant barrier for the introduction of macromolecules into Chlamydomonas cells (Jeon et al., 2017; this work). Indeed, when precise editing of walled strains is attempted by co-electroporation of a CRISPR/Cas9 RNP, a PCR fragment or a plasmid encoding a transgene expressing an antibiotic resistance gene (for selection of the small fraction of cells taking up the editing components) and a donor DNA (to be used as homologous template for HDR), most Chlamydomonas cells appear to repair the induced DSBs by NHEJ pathways, based on the sequence analyses of recovered colonies (Greiner et al., 2017; this work). Such repair often involves the undesirable incorporation of intact or frequently jumbled antibiotic resistance gene sequences at the site of Cas9 DNA cleavage (Greiner et al., 2017; this work).

Given these constraints, we tested whether co-targeting two genes by electroporation of CRISPR/Cas9 RNPs and template ssODN donors would facilitate the recovery of precise edits in a gene of interest after selecting for precise editing of the other gene (corresponding to a selectable marker). Conceptually, this strategy is expected to select for cells capable of taking up the editing components and expressing the machinery required for HDR after transfection. We first chose *PPX1*, encoding protoporphyrinogen oxidase, as a selectable marker since a single base pair change (G->A) within the protein coding sequence confers resistance to porphyric herbicides such as oxyfluorfen (Randolph-Anderson *et al*., 1998; Brueggeman *et al*., 2014). Selecting electroporated cells on oxyfluorfen containing medium allowed the recovery of colonies edited in two co-targeted genes, namely *FTSY* and *WDTC1*. On average, between 2 and 4% of oxyfluorfen resistant colonies showed sequence alterations in the co-targeted genes, and about one third of these colonies showed precise, scarless editing mediated by HDR. These observations suggest that our co-editing approach does facilitate the isolation of colonies precisely edited in genes of interest in walled Chlamydomonas. Moreover, the co-editing approach avoids the incorporation of exogenous selectable marker gene fragments into CRISPR/Cas9 cleavage sites, which makes it difficult to obtain scarless gene edits, as observed in previous gene editing protocols (Shin et al., 2016; Greiner et al., 2017; Kang et al., 2020; Picariello et al., 2020) and once more documented in the present study (Fig. 1 and Supplemental Fig. 2).

The heart of the scheme presented here is to expedite the recovery of a specific mutation in any Chlamydomonas gene, in the absence of a selectable phenotype for that mutation. Selection of the desired mutant is based on the ability to simultaneously deliver into cells two pairs of components: a CRISPR/Cas9 RNP and ssODN donor designed to modify the target gene of interest along with a second pair consisting of CRISPR/Cas9 (*PPX1*) RNP and a ssODN donor designed to convert the *PPX1* gene into a selectable marker gene. Our isolation of oxyfluorfen resistant colonies containing precise, scarless mutations in the *FTSY* and *WDTC1* genes is proof of concept for this approach. Because each additional mutation that an investigator wishes to incorporate into a strain requires another gene that can be converted to a selectable marker, we also demonstrated that the *ALS* gene could serve as an additional selectable marker. Preliminary experiments indicate that the argininosuccinate lyase (*ARG7*) gene, of the arginine auxotroph *arg7-8*, could also be used as a selectable marker since conversion to the wild-type sequence can create easily selected autotrophic cells (Jiang and Weeks, 2017; Jiang et al., 2017). Other endogenous genes could potentially be edited by single base pair substitutions to become selectable markers, such as those encoding phytoene desaturase (PDS), conferring resistance to the bleaching herbicide norflurazon (Brueggeman et al., 2014; Suarez et al., 2014), and cytosolic ribosomal protein S14 (CRY1), conferring resistance to the antibiotics cryptopleurine and emetine (Nelson et al., 1994; Neupert et al., 2009). These examples of endogenous genes with demonstrated or potential utility as selectable markers suggest it may soon be possible to attempt the creation of Chlamydomonas cell lines containing precise mutations in multiple target genes, directed at aiding the study of complex cellular functions or metabolic pathways of academic or biotechnological interest. We also note that our strategy is ‘transgene-free’ since, in the instances of precise gene editing, exogenous DNA is not expected to be incorporated into the genome, although potential off-target effects mediated by CRISPR/Cas9 RNPs remain to be examined.

If the presence of transgenes is not a concern, the Hegemann laboratory (Sizova et al. 2013; Greiner et al., 2017) demonstrated the ability to transform into the genome of Chlamydomonas an antibiotic resistance gene (i.e., an *aphVIII* transgene) that had been deliberately inactivated by a simple sequence modification. Using this transgenic strain, they subsequently employed CRISPR/Cas9-mediated editing to modify the mutant *aphVIII* gene and restore antibiotic resistance. A similar approach, using inactivated versions of the many antibiotic resistance genes effective in Chlamydomonas, has the potential to greatly broaden the spectrum of editable genes available for selection purposes in co-targeting experiments.

Sequence analyses of the recovered Chlamydomonas colonies provided insights regarding the mechanism of DSB repair involving ssODN templates. As described in several eukaryotes (Gallagher and Haber, 2017; Kan et al., 2017; Paix et al., 2017; Boel et al., 2018; Harmsen et al., 2018; Richardson et al., 2018; Sansbury et al., 2019; Gallagher et al., 2020), the single-strand template repair (SSTR) model appears to explain best the repair of Cas9-induced DSBs with ssODNs as homologous donor DNA. In SSTR, ssODNs are used for the synthesis of complementary DNA, rather than being integrated into the genome, and the process is polarity sensitive and dependent, unlike other homologous repair mechanisms, on the Fanconi anemia pathway (Gallagher and Haber, 2017; Kan et al., 2017; Paix et al., 2017; Boel et al., 2018; Harmsen et al., 2018; Richardson et al., 2018). As shown diagrammatically in Supplemental Fig. 1, for the proposed repair of *FTSY*, a DSB generated by Cas9 is expected to be resected to yield 3’ overhangs on both sides of the DSB. However, only one of the 3’ overhangs can pair with the homologous ssODN donor, which confers polarity to the repair, and primes the synthesis of a complementary DNA strand. Bridging of the DSB is eventually accomplished when the newly synthesized strand is displaced from the ssODN donor and anneals with the complementary strand at the locus. Finally, DNA polymerases and ligases complete the DSB repair. As proposed in mammalian cells (Kan et al., 2017; Harmsen et al., 2018), the likely erosion of the ends of 3’ overhangs would allow the introduction of edits very close to the DSB by *de novo* synthesis of both DNA strands (i.e., by gap filling) (Supplemental Fig. 1), avoiding the formation of heteroduplex DNA and interference from DNA mismatch repair mechanisms.

In higher plants, ssODNs have been used for precise gene editing, as templates for the repair of DSBs generated by CRISPR/Cas9, but very few edited plants/calli have been analyzed by sequencing precluding any inference on the repair mechanism (Shan et al., 2013; Svitashev et al., 2015; Sauer et al., 2016). In the moss *Physcomitrella patens*, it has been suggested that a template (ssODN)-target pairing mechanism is involved in the repair of DSBs generated by CRISPR/Cas9 (Yi and Goshima, 2020). In our work with Chlamydomonas, several lines of evidence, taken together, support the interpretation that SSTR is responsible for HDR. First, synthesis of a DNA strand complementary to a ssODN donor begins at the DSB and, since repair DNA polymerases are not very processive (Rodgers and McVey, 2016), nucleotide changes closer to the DSB are more likely to be incorporated into the genome (Paquet et al., 2016; Kan et al., 2017; Paix et al., 2017; Boel et al., 2018). Additionally, since the locus strand complementary to the newly synthesized strand may often be somewhat eroded at its 3’ end (Dorsett et al., 2014; Harmsen et al., 2018), the generation of heteroduplex DNA (which could be corrected back to the wild type sequence by DNA mismatch repair mechanisms) may be avoided closer to the DSB. These predictions are consistent with the greater incorporation efficiency of G->A (closer to the DSB) vs. C->T (more distant from the DSB) in edited *PPX1* (Fig. 2C). Second, SSTR is unidirectional since only one of the 3’ overhangs at the DSB can pair with the ssODN donor to synthesize a complementary strand (Gallagher and Haber, 2017; Paix et al., 2017; Boel et al., 2018). In Chlamydomonas repair polarity is best supported by the analysis of colonies where one side of the DSB appears to show integration of donor DNA by homologous recombination (i.e., a crossover) and the other side by NHEJ. Close inspection of the sequences at the target site in these colonies (for both *FTSY* and *WDTC1*) indicates strict polarity (Supplemental Fig. 2, colony 13b; Supplemental Fig. 5, colony 4-1; Supplemental Fig. 7, colony 2A-10; Supplemental Fig. 8, colony 63). The DSB side with apparent integration by a crossover event is always the side that would prime DNA synthesis using the ssODN donor as template. A more plausible explanation is that SSTR started correctly, with the synthesis of a DNA strand complementary to the ssODN donor, but the repair was eventually resolved by NHEJ.

Third, repair DNA polymerases, such as those involved in the synthesis of a strand complementary to the ssODN donor, are error prone and cause base pair substitutions and frameshift mutations (Rodgers and McVey, 2016; Gallagher and Haber, 2017; Richardson et al., 2018; Gallaher et al., 2020). This is consistent with the recovery of several Chlamydomonas colonies where the intended nucleotide modifications in the *FTSY* or *WDTC1* genes did occur, but additional sequence changes typical of replicative errors (such as single base pair deletions or substitutions) were also observed (Fig. 3C, colony 3-2; Fig. 4C, colony 2B-19). Similar findings have already been reported for CRISPR/Cas12a editing with ssODN templates in cell-wall-less Chlamydomonas strains (Ferenczi et al., 2017). Fourth, ssODN-directed repair is prone to template switching between donor DNA molecules, an event apparently dependent on regions of microhomology (Rodgers and McVey, 2016; Paix et al., 2017; Boel et al., 2018). This appears to have happened at least in one case in Chlamydomonas, partly copying two molecules of *WDTC1* ssODN donor DNA (Supplemental Fig. 6B and Supplemental Fig. 7, colony 1B-24).

In summary, our co-editing approach allows the recovery of precise sequence edits on genes of interest at a practical frequency, often requiring the PCR analysis of little more than 100 herbicide resistant colonies (at least for the genes currently tested). Moreover, since we used walled Chlamydomonas cells, quite refractory to CRISPR/Cas RNP transfection (Ferenczi et al., 2017; Picariello et al., 2020; this work), we anticipate that removal of the cell wall by autolysin treatment (prior to electroporation) may improve the efficiency of our protocol, as clearly demonstrated for targeted insertional mutagenesis (Picariello et al., 2020). However, the current strategy will largely be useful for the introduction of relatively short edits, for instance to investigate the function of specific amino acid residues in proteins expressed in their endogenous context. In mammalian cells it has been proposed that Cas9-stimulated repair using ssODN donors mimics some substrate of the Fanconi anemia pathway, such as a stalled replication fork (Richardson et al., 2018). This diverts DSB repair through SSTR, which is effective but has the drawback of being prone to replicative errors, as also appears to be the case in Chlamydomonas.

## Materials and Methods

### Strains, culture conditions and lipid accumulation

*C. reinhardtii* strains g1 (CC-5415; *nit1*, *agg1*, *mt+*), CC-124 (*nit1*, *nit2*, *agg1*, *mt-*) (Chlamydomonas Resource Center, https://www.chlamycollection.org) or derived edited mutants were used in all reported experiments. Unless noted otherwise, cultures were incubated under continuous illumination (150 µmol m^−2^ s^−1^ photosynthetically active radiation) on an orbital shaker (190 rpm) at 25 °C and ambient CO_2_ levels. For transfection experiments, cells were precultured, after inoculation from one-week-old plates, to an optical density of ∼0.4 at 750 nm in Tris-Acetate-Phosphate (TAP) medium (Harris, 1989) supplemented with 1 μg ml^−1^ cyanocobalamin (vitamin B_12_), to enhance Chlamydomonas thermal tolerance (Xie et al., 2013). A one-tenth aliquot was then transferred into fresh medium and grown to middle logarithmic phase (∼1 to 2 x 10^6^ cells ml^−1^). Prior to electroporation, cells were collected by centrifugation and resuspended in TAP medium containing 40 mM sucrose and 1 μg ml^−1^ cyanocobalamin to a final density of ∼2.2 x 10^8^ cells ml^−1^. For growth experiments, cells were pre-cultured as described before and then inoculated into minimal (Sueoka, 1960) medium (phototrophic conditions) or TAP medium (mixotrophic conditions). Culture growth was examined by measuring daily absorbance at 750 nm. For lipid accumulation analyses, cells were inoculated into TAP medium lacking nitrogen and 200 μl-aliquots were tranferred daily to a multi-well plate and mixed with Nile Red (Sigma, 72485) to a final concentration of 1 μg ml^−1^ (Msanne et al., 2012). Nile Red fluorescence (excitation at 488 nm; emission at 565 nm) was measured in a multi-well plate reader (Synergy H1, Biotek), normalized to cell density (determined as absorbance at 750 nm) and expressed in arbitrary units (Kim et al., 2018).

### crRNA, tracrRNA, ssODN donors and *aphVIII* transgenic DNA

The 19- or 20-nt guide sequences, corresponding to the target-specific protospacer regions (Supplemental Table 7), were designed using the Cas-Designer (www.rgenome.net/cas-designer) or Chopchop (https://chopchop.cbu.uib.no) websites. Each CRISPR RNA (crRNA) was synthesized as a custom, chemically-modified 35- or 36-nt oligo (containing 16 additional, common nucleotides for annealing to the tracrRNA) by Integrated DNA Technologies (IDT). We used a commercially available trans-acting CRISPR RNA (tracrRNA) (IDT, 1072534 Alt-R^®^ CRISPR-Cas9 tracrRNA). Single-stranded oligodeoxynucleotide donors (Supplemental Table 2) were designed overlapping the CRISPR/Cas9 cleavage site, but with different lengths of homology arms (from 30 to 70 nucleotides), and synthesized as Ultramer DNA Oligos (IDT). The paromomycin resistance cassette (*aphVIII* transgene) was amplified by PCR from the pSI103 plasmid (Sizova et al., 2001).

### Preparation of CRISPR/Cas9 RNPs

CRISPR/Cas9 crRNA and tracrRNA (each at ∼53 μM final concentration) in 1.1x NEB 3.1 buffer (New England Biolabs, B7203S) were annealed by placing a microcentrifuge tube in a beaker with water heated to 96 °C and then allowed to cool slowly to room temperature. To assemble the RNP complex, one volume of Alt-R^®^ S.p.Cas9 Nuclease V3 (IDT, 1081059) was mixed with 3.5 volumes of annealed crRNA/tracrRNA and incubated at 37 °C for 20 min. In this mixture, the guide RNA is present at ∼3-fold molar excess relative to the Cas9 protein and, if saturation binding is achieved, the RNP final concentration would be ∼13.6 μM.

### Chlamydomonas transfection

The initial experiments (Supplemental Tables 1 and 3) were carried out following the method of Greiner et al. (2017). In order to improve the editing frequency, we subsequently introduced several modifications that resulted in an optimized protocol. For single gene editing experiments, 2.3 μl of the CRISPR/Cas9 (*FTSY*) RNP [or the CRISPR/Cas9 (*WDTC1*) RNP] were mixed with 1.5 μl of the PCR product (∼1.0 μg of dsDNA in TE buffer) encoding the *aphVIII* transgene (± 225 pmol of the corresponding ssODN donor). For co-editing experiments, the CRISPR/Cas9 (*PPX1*) RNP and the CRISPR/Cas9 (*FTSY*) RNP or the CRISPR/Cas9 (*WDTC1*) RNP were first mixed in a 1 to 3 ratio (i.e., a 3-fold molar excess of the RNP targeting the unselected gene of interest). Three microliters of the combined RNPs were then added to 1.5 μl of pre-mixed (in TE buffer) *PPX1* (at 50 μM) and *FTSY* (or *WDTC1*) (at 150 μM) ssODN donors. For each electroporation, an aliquot of 36 μl of cells (∼7.9 x 10^6^ cells ml^−1^), resupended in TAP medium containing 40 mM sucrose and 1 μg ml^−1^ cyanocobalamin, was mixed with 3.8 μl of the RNP/*aphVIII* dsDNA/ssODN or 4.5 μl of the combined RNPs/ssODNs and placed in an electroporation cuvette with a 2 mm gap. Transfection was performed with a NEPA21 electroporator (Nepa Gene Co.), with adjusted parameters relative to a published protocol (Yamano et al., 2013). Prior to elecroporation the impedance was adjusted to 0.25-0.28 kΩ by adding more resuspended cells or withdrawing from the mixture in the cuvette (stepwise, 3 μl at a time). Electroporation was carried out by using two 6-ms/250-V poring pulses at 50-ms intervals and a decay rate of 40%, followed by five 50-ms/20-V polarity-exchanged transfer pulses at 50-ms intervals and decay rate of 40%. Immediately after electroporation, the cells were diluted in 500 μl of TAP medium containing 40 mM sucrose and 1 μg ml^−1^ cyanocobalamin and transferred to a 1.5-ml microcentrifuge tube.

### Post-electroporation heat shock, cell recovery and plating

Electroporated cells were placed on an orbital shaker (50 rpm) under dim lights for 3 hours at room temperature. Cells were then heat shocked at 39 °C for 30 min, with gentle agitation, in a water bath. Subsequently, cells were incubated again on the orbital shaker for ∼40 hours. After completion of this recovery period, cells from each electroporation were spread on two TAP agar plates containing the appropriate selective agent (12 μg ml^−1^ paromomycin or 0.18 μM oxyfluorfen). The plates were incubated at room temperature under continuous light (∼100 µmol m^−2^ s^−1^ photosynthetically active radiation) until visible colonies appeared (usually about two weeks).

### Testing the acetolactate synthase (*ALS1*) gene as a selectable marker

Cells of the wild-type strain CC-124 were grown in TAP medium, in 5% CO_2_ at 25 °C with shaking and continuous light, to mid-log phase and then concentrated to ∼5 x 10^7^ cells ml^−1^ in TAP medium containing 60 mM sucrose. A mixture consisting of ∼100 pmol Cas9 (New England Biolabs, M0386M) and ∼850 pmol of *ALS1* crRNA (Supplemental Table 7) preannealed with an equimolar amount of tracrRNA, in a total volume of 30 μl in NEB Cas9 buffer, was allowed to incubate at 37 °C for 30 min. Then, 1μl (∼800 pmol) of the *ALS1* ssODN donor (Supplemental Table 2) was added, just before cell transfection. This mixture was added to 0.5 ml of concentrated cells (∼2.5 x 10^7^ cells) and placed in an electroporation cuvette with a 4 mm gap for 5 min at 16 °C. Electroporation was carried out with a Gene Pulser Xcell system (Bio-Rad) set at 750 V, 25 μF and ∞ resistance (to produce a time constant of 5 to 6 ms). After electroporation, cells were allowed to rest for 10 min and then diluted into 50 ml of TAP containing 60 mM sucrose and incubated for 24 hours in 5% CO_2_ at 25 °C with shaking and light. No post-electroporation heat shock treatment was applied. Cells were eventually concentrated by centrifugation, resuspended in ∼0.2 ml TAP, split into two aliquots and spread on TAP agar plates containing sulfometuron methyl (SMM) at 5 μM. After incubation for ∼14 days in 5% CO_2_ at 25°C in continuous light, colonies were picked and restreaked on TAP + SMM plates to isolate single colonies. DNA was extracted from each colony and used for PCR amplification and DNA sequencing of the *ALS1* target site.

### Genotyping of potentially edited mutants

When targeting the *PPX1* gene, a subset of colonies surviving on oxyfluorfen containing medium was examined by colony PCR amplification (Cao et al., 2009) of the target site followed by sequencing of the PCR products. When targeting the *FTSY* gene, only pale-green colonies surviving on paromomycin containing medium or oxyfluorfen containing medium (in case of co-editing with the *PPX1* gene) were examined by colony PCR amplification of the target site. All detectable PCR products, those of expected size and those differing in size from the wild-type amplicon (due to possible insertions or deletions), were examined by sequencing. PCR products of the expected size were also characterized by digestion with the *Nhe*I enzyme (Sambrook and Russell, 2001). When targeting the *WDTC1* gene, colonies surviving on paromomycin containing medium or oxyfluorfen containing medium (in case of co-editing with the *PPX1* gene) were examined by colony PCR amplification of the target site followed by digestion of the PCR product with the *Bss*HII enzyme (Sambrook and Russell, 2001). Only PCR products successfully digested with *Bss*HII or potentially having indels were characterized by sequencing. In all cases, amplification of the Actin gene was used as a positive control. To ensure amplification of the correct DNA fragments from the target genes, in most cases nested polymerase chain reactions were carried out with the primers listed in Supplemental Table 8. The PCR conditions for general amplification were 35 cycles at 94 °C for 30 s, at 50 °C for 30 s and at 71 °C for 90 s. Aliquots (8 µl) of each PCR were resolved on 1.5% agarose gels and visualized by ethidium bromide staining (Sambrook and Russell, 2001).

### Semi-quantitative reverse transcriptase (RT)-PCR analyses of *WDTC1* transcript abundance

Total cell RNA, from cells harvested in the middle of the logarithmic phase, was purified with TRI Reagent (Molecular Research Center), following the manufacturer’s instructions. Reverse transcription reactions were performed as previously described (Carninci et al., 1998) using Superscript^TM^ III (Invitrogen, 18980051). The synthesized cDNA was then used as a template in standard PCRs (Sambrook and Russell, 2001) using a number of cycles showing a linear relationship between input RNA and the final product, as determined in preliminary experiments. The PCR conditions for amplification of the Actin control were 25 cycles at 94 °C for 30 s, at 58 °C for 30 s and at 71 °C for 30 s. The PCR conditions for amplification of the *WDTC1* transcript were 30 cycles at 94 °C for 30 s, at 60 °C for 30 s and at 71 °C for 30 s. Aliquots (8 µL) of each RT-PCR were resolved on 1.5% agarose gels and visualized by ethidium bromide staining (Sambrook and Russell, 2001). All primers used for RT-PCR are listed in Supplemental Table 8.

### Examination of CRISPR/Cas9 (*FTSY*) RNP cellular uptake by fluorescence microscopy

Chlamydomonas cells were electroporated with a commercially available trans-acting CRISPR RNA conjugated to the ATTO 550 fluorophore (IDT, 1075928 Alt-R® CRISPR-Cas9 tracrRNA, ATTO™ 550), either alone or assembled into a CRISPR/Cas9 (*FTSY*) RNP (using ∼2-fold molar excess of the Cas9 protein relative to the tracrRNA-ATTO 550). Cell were examined for uptake of the transfected macromolecules at 1, 4, and 24 hours after electroporation. However, the reported data corresponds to 4 h after electroporation, when the strongest signal was detected. For microscopy analyses, cells from a single electroporation were pelleted by centrifugation, washed twice and eventually resuspended in TAP medium containing 40 mM sucrose, to which a 1/100 volume of iodine (1% in ethanol) was added. Cells were visualized with an EVOS FL Auto Cell Imaging System (ThermoFisher Scientific), using the RFP light cube (Excitation 542/20, Emission 593/40) for ATTO 550 fluorescence and the Cy5 light cube (Excitation 635/18, Emission 692/40) for chlorophyll fluorescence. Pseudo-colors were used to represent signals from the different channels.

## Supplemental Data

The following supplemental materials are available.

**Supplemental Figure 1.** Single-strand template repair (SSTR) model for homology directed repair of the *FTSY* gene, using as complementary template the transfected ssODN.

**Supplemental Figure 2.** DNA sequences of *FTSY* disrupted mutants obtained by co-transfection of CRISPR/Cas9 (*FTSY*) RNP, a dsDNA PCR product encoding the *aphVIII* transgene and the *FTSY* ssODN donor.

**Supplemental Figure 3.** Fluorescence microscopy analysis of the cellular uptake of the CRISPR/Cas9 (*FTSY*) RNP after electroporation.

**Supplemental Figure 4.** Single-strand template repair (SSTR) model for homology directed repair of the *PPX1* gene, using as complementary template the transfected ssODN.

**Supplemental Figure 5.** DNA sequences of *FTSY* insertional mutants obtained by co-targeting the *PPX1* and *FTSY* genes for CRISPR/Cas9 editing.

**Supplemental Figure 6.** Single-strand template repair (SSTR) model for homology directed repair of the *WDTC1* gene, using as complementary template the transfected ssODN.

**Supplemental Figure 7.** DNA sequences of *WDTC1* insertional mutants obtained by co-targeting the *PPX1* and *WDTC1* genes for CRISPR/Cas9 editing.

**Supplemental Figure 8.** DNA sequences of *WDTC1* insertional mutants obtained by co-transfection of CRISPR/Cas9 (*WDTC1*) RNP, a dsDNA PCR product encoding the *aphVIII* transgene and the *WDTC1* ssODN donor.

**Supplemental Figure 9.** Single-strand template repair (SSTR) model for homology directed repair of the *ALS1* gene, using as complementary template the transfected ssODN.

**Supplemental Figure 10.** DNA sequence of the *ALS1* target site in sulfometuron methyl resistant colonies.

**Supplemental Table 1.** Efficiency of *FTSY* targeted gene disruption by electroporation with CRISPR/Cas9 (*FTSY*) RNP and a PCR product encoding the *aphVIII* transgene.

**Supplemental Table 2.** Single-stranded oligodeoxynucleotides (ssODNs) electroporated into cells as templates for HDR.

**Supplemental Table 3.** Efficiency of *PPX1* gene editing by electroporation with CRISPR/Cas9 (*PPX1*) RNP and ssODN donor DNA.

**Supplemental Table 4.** Efficiency of *FTSY* gene editing by co-electroporation with CRISPR/Cas9 (*FTSY*) RNP, CRISPR/Cas9 (*PPX1*) RNP and the corresponding ssODN donor DNAs.

**Supplemental Table 5.** Efficiency of *WDTC1* gene editing by co-electroporation with CRISPR/Cas9 (*WDTC1*) RNP, CRISPR/Cas9 (*PPX1*) RNP and the corresponding ssODN donor DNAs.

**Supplemental Table 6.** Mutagenesis of the Chlamydomonas acetolactate synthase (*ALS1*) gene using CRISPR/Cas9 (*ALS1*) RNP and *ALS1* ssODN donor DNA.

**Supplemental Table 7.** CRISPR RNAs (crRNAs) used in the study.

**Supplemental Table 8.** PCR primers used in the study.

## Acknowledgments

This work was supported in part by grants from the National Science Foundation (MCB 1616863) and the Gordon and Betty Moore Foundation (Award No. 4968.01) to H.C. We thank Samantha Rau and Rakim Ali for technical assistance with the colony PCR analyses.

